# PSYLLIUM FIBER PROTECTS MICE AGAINST WESTERN DIET-INDUCED METABOLIC SYNDROME VIA MICROBIOTA-DEPENDENT MECHANISM

**DOI:** 10.1101/2022.12.15.520605

**Authors:** Alexis Bretin, Andrew T. Gewirtz

## Abstract

We sought to identify semi-purified fibers that broadly promote gut and metabolic health and, moreover, identify the mechanisms by which they do so. Screening an array of purified fibers led us o appreciate that, of all fibers tested, psyllium was unique in that it strongly protected mice against both colitis and metabolic syndrome. Further investigation led us to appreciate that psyllium’s mechanism of action is quite distinct. Specifically, data we report in a manuscript under review indicates that low doses of psyllium protect mice against colitis via increasing serum bile acids that activate FXR. Here, we report that higher doses of psyllium protect mice against diet-induced obesity/metabolic syndrome by a FXR-independent mechanism that requires presence of at least a minimal microbiota.

## Introduction

Our published work has shown that the fermentable fiber psyllium protects against diet-induced metabolic syndrome ^1^. However, inulin makes mice highly prone to experimentally induced colitis, especially in the DSS model ^2^. Hence, we have been screening other fibers seeking one tht would protect against both colitis and metabolic syndrome. A manuscript that was recently positively reviewed reports that psyllium protects against colitis via mechanism dependent upon the FXR bile acid receptor ^3^. This manuscript reports psyllium’s impacts on metabolic syndrome. A full manuscript will be submitted for peer review in the future.

## Methods

### Mice

C57BL/6 WT and FXR^-/-^ mice were purchased from Jackson Laboratory (Bar Harbor, ME). Gnotobiotic C57BL/6 germ-free mice (GF) were purchased from Taconic Biosciences Inc, (Rensselaer, NY). Mice harboring Altered Schaedler Flora (ASF) were generated from these GF mice as previously described ^4^. All mice were bred, maintained, and experimentally studied at Georgia State University under an institutional animal care and use committee (IACUC protocol # A17047).

### Diets and HFD-induced metabolic syndrome

Mice, 6–8 weeks of age, were fed either a standard grain-based chow (GBC, Purina 5001) or a purified “open-source”, herein referred to as compositionally-defined, diet (CDD), the composition of which is indicated in Supplementary Table 1 and fed *ad libitum* for 4 weeks. Mice were monitored weekly for weight gain. At day 28, mice were euthanized, serum, colon length, washed colon weight, epididymal fat pad, cecum and spleen weights were collected for downstream analysis.

### Glucose measurement

Glucose tolerance was measured 25 days following initiation of indicated diet. Mice were placed in a clean cage and provided with water but without food for 5 hours. Baseline blood glucose was then measured using a Nova Max plus Glucose meter. Mice were administered glucose, 2 mg of glucose/gm body weight, and blood glucose levels measured 30, 60, and 90 minutes later. Data are expressed as mg glucose/dL blood. 2 days later, (i.e. day 27), to measure “fasting glucose levels”, mice were placed in a clean cage and provided with water but without food for 5h, at which point, fasting glucose levels were determined and insulin tolerance test initiated. Specifically, 5h-fasted mice were injected with 0.5 U insulin/kg body weight and blood glucose levels were measured at 30, 60, 90 min after injection.

### RNA Extraction and Real-Time PCR

Total RNA was isolated from colon using TRIzol (Invitrogen, Carlsbad, CA); the expression level of IL-6, CXCL1, and TNF-α was analyzed by using quantitative real-time PCR according to the Biorad iScript One-Step RT-PCR Kit in a CFX96 apparatus (Bio-Rad, Hercules, CA) with the following primers (F/R): IL-6: GTGGCTAAGGACCAAGACC, GGTTTGCCGAGTAGACCTCA; CXCL1: TTGTGCGAAAAGAAGTGCAG, TACAAACACAGCCTCCCACA; TNF-a: CGAGTGACAAGCCTGTAGCC, CATGCCGTTGGCCAGGA; 36B4: TCCAGGCTTTGGGCATCA, CTTTATTCAGCTGCACATCACTCAGA. Differences in transcript levels were quantified by normalization of each amplicon to housekeeping gene 36B4.

### Bacterial Quantification in Feces

To measure the total fecal bacterial load, total DNA was isolated from weighted feces using QIAamp DNA Stool MiniKit (Qiagen, Hilden, Germany). DNA was then subjected to qPCR using QuantiFast SYBR Green PCR kit (Bio-Rad, Hercules, CA) with universal 16S rRNA primers 8F: 50-AGAGTTTGATCCTGGCTCAG-30 and 338R: 50-CTGCTGCCTCCCGTAGGAGT-30 to measure total bacteria number. Results are expressed as bacteria number per mg of stool using a standard curve.

### Short chain fatty acids (SCFA) quantification

#### Statistical Analyses

Statistical significances of results were analyzed by analysis of variance (ANOVA test). Significance is express as *P < 0.05, **P < 0.01, ***P < 0.001, ****P < 0.0001.

## RESULTS

Groups of mice that had been maintained on grain-based chow (GBC) were maintained on this diet or switched to a compositionally-defined low-fat diet (LFD), an obesogenic high-fat diet (HFD), which contained 50g/kg fiber, namely cellulose or HFD enriched with 100 g/kg cellulose (Cell), inulin (Inul), pectin (Pec), psyllium (Psy), or Hi-Maize (Hi-M). As expected mice fed the low-fiber obesogenic HFD, exhibited a stark increase in adiposity, which was accompanied by dysglycemia as reflected by measure of glucose and insulin tolerance. As expected, fermentable fibers inulin and pectin protected against these indices of metabolic syndrome. Yet, enrichment of HFD with psyllium provided even greater protection in this disease model, while Hi-Maize had minimal impacts (Figure 1). HFD-induced metabolic syndrome associates with loss of colon and cecal mass, both of which were also restored by enriching HFD with psyllium. Thus, psyllium appeared to meet our goal of a fiber that could protect against colitis and metabolic syndrome.

**Figure 1:**
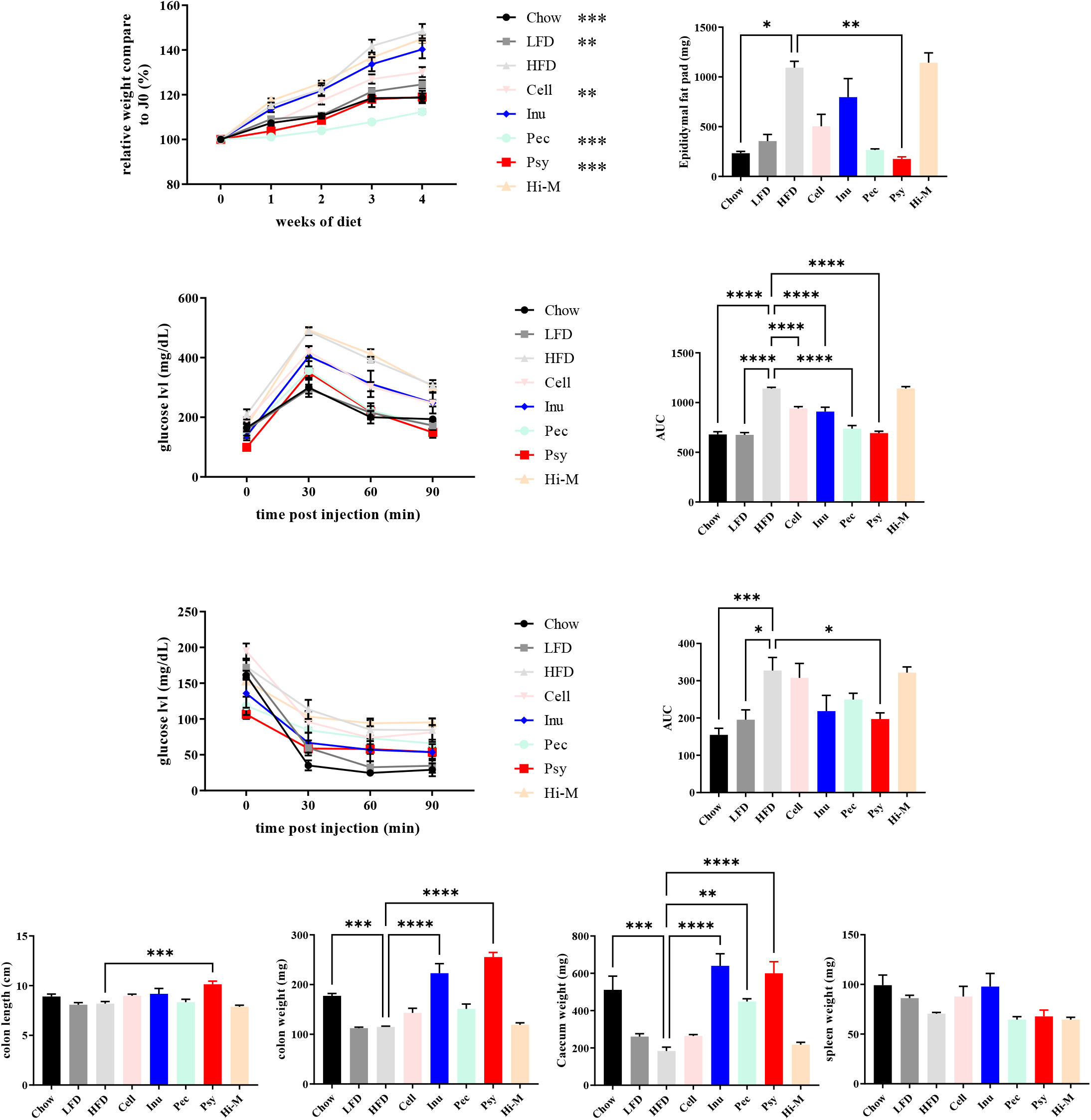
Psyllium provides the strongest protection against HFD-induced Metabolic syndrome. Male 6-8 week-old C57Bl/6 mice were fed with the indicated diet for 28 day. **A**. Relative body weight over time. **B**. Epididymal fat pad weight. **C**. Glucose tolerance test and respective areas under the curve. **D**. Insulin tolerance test and respective areas under the curve. **E**. Colon length. **D**. Colon weight. **E**. Cecum weight. **F**. Spleen weight. Data are expressed as means ± SEM of N=5 mice per group and are representative of 3 independent experiments. Significance was determined by ANOVA variance test. *P < 0.05, **P < 0.01, ***P < 0.001, ****P < 0.0001.

We next investigated the dose responsiveness of psyllium’s impact in diet-induced obesity (Figure 2). A significant reduction in diet-induced adiposity was observed with diets comprised of as little as 4% psyllium by weight while a dose of 8% reduced adiposity to levels similar to GBC-fed mice. Reductions in adiposity from various doses of psyllium were generally associated with improved glycemic control and restoration of intestinal mass.

**Figure 2:**
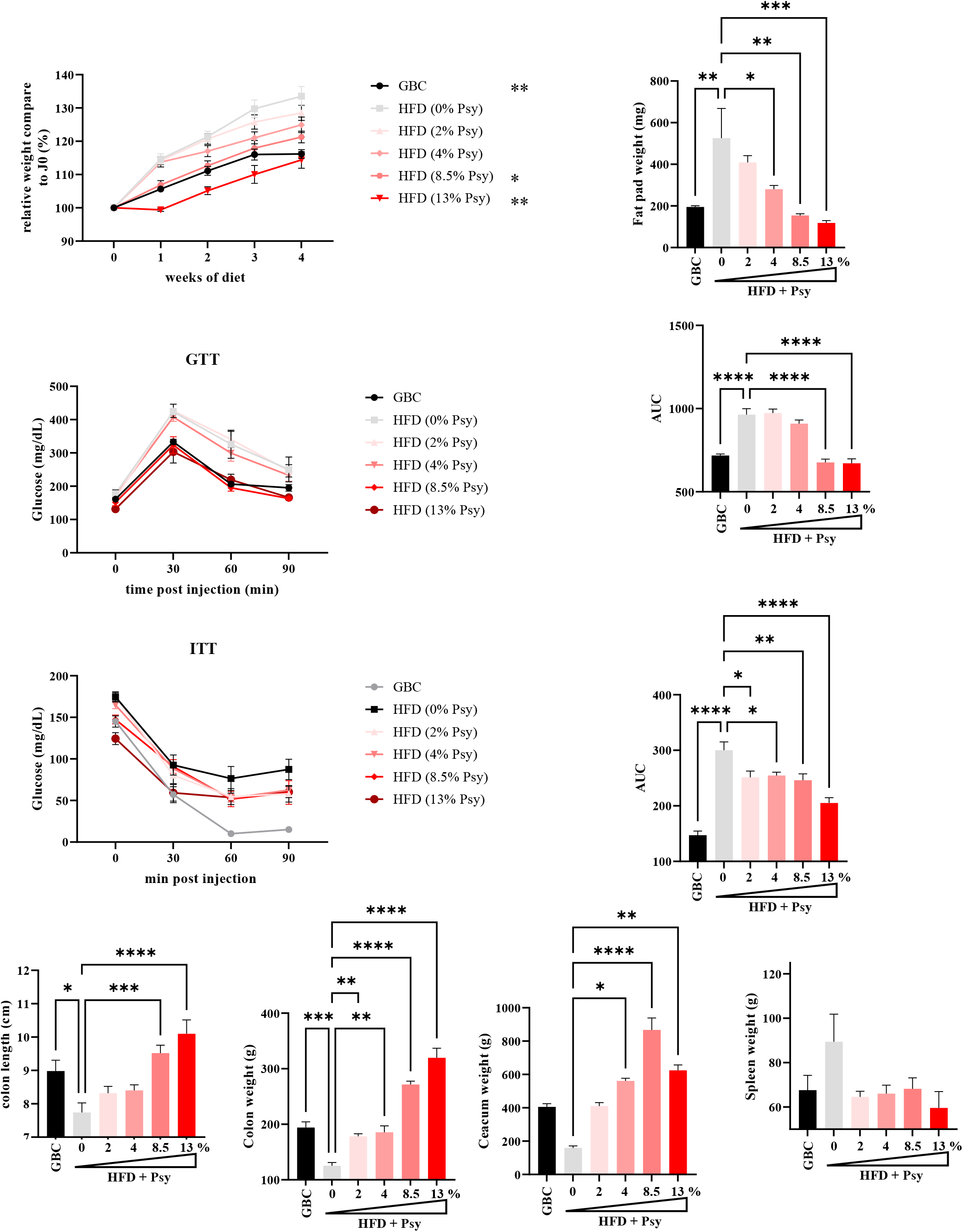
Psyllium protects against HFD-induced metabolic syndrome in a dose-dependent manner with an optimum protection over 8.5% psyllium. Male 6-8 week-old C57Bl/6 mice were fed with the indicated diet for 28 day. **A**. Relative body weight over time. **B**. Epididymal fat pad weight. **C**. Glucose tolerance test and respective areas under the curve. **D**. Insulin tolerance test and respective areas under the curve. **E**. Colon length. **D**. Colon weight. **E**. Cecum weight. **F**. Spleen weight. Data are expressed as means ± SEM of N=5 mice per group. Significance was determined by ANOVA variance test. *P < 0.05, **P < 0.01, ***P < 0.001, ****P < 0.0001.

We next turned our attention to investigating the mechanism by which psyllium protected against metabolic syndrome. Psyllium’s protection against colitis requires the FXR bile acid receptor. We reasoned that its protection against diet-induced obesity may also utilize this mechanism. However, psyllium’s protection against HFD-induced in FXR-KO mice arguing against this possibility (Figure 3). Psyllium is viewed as a semi-soluble partially fermentable fiber suggesting that fermentation contributed to its impacts. However, psyllium did not result in an increase in short-chain fatty acids (Figure 4). Nor was psyllium’s protection against HFD-induced metabolic syndrome prevented by blocking fermentation with hops B-acids. These findings argue against a role for fermentation. Inulin elicits copious amounts of IL-22, which is required for inulin’s beneficial impacts on metabolic syndrome. In contrast, psyllium lowered IL-22 levels in the gut arguing against a mechanism analogous to inulin. In further contrast to inulin, which increased gut bacterial density, psyllium reduced this parameter (Figure 5).

**Figure 3:**
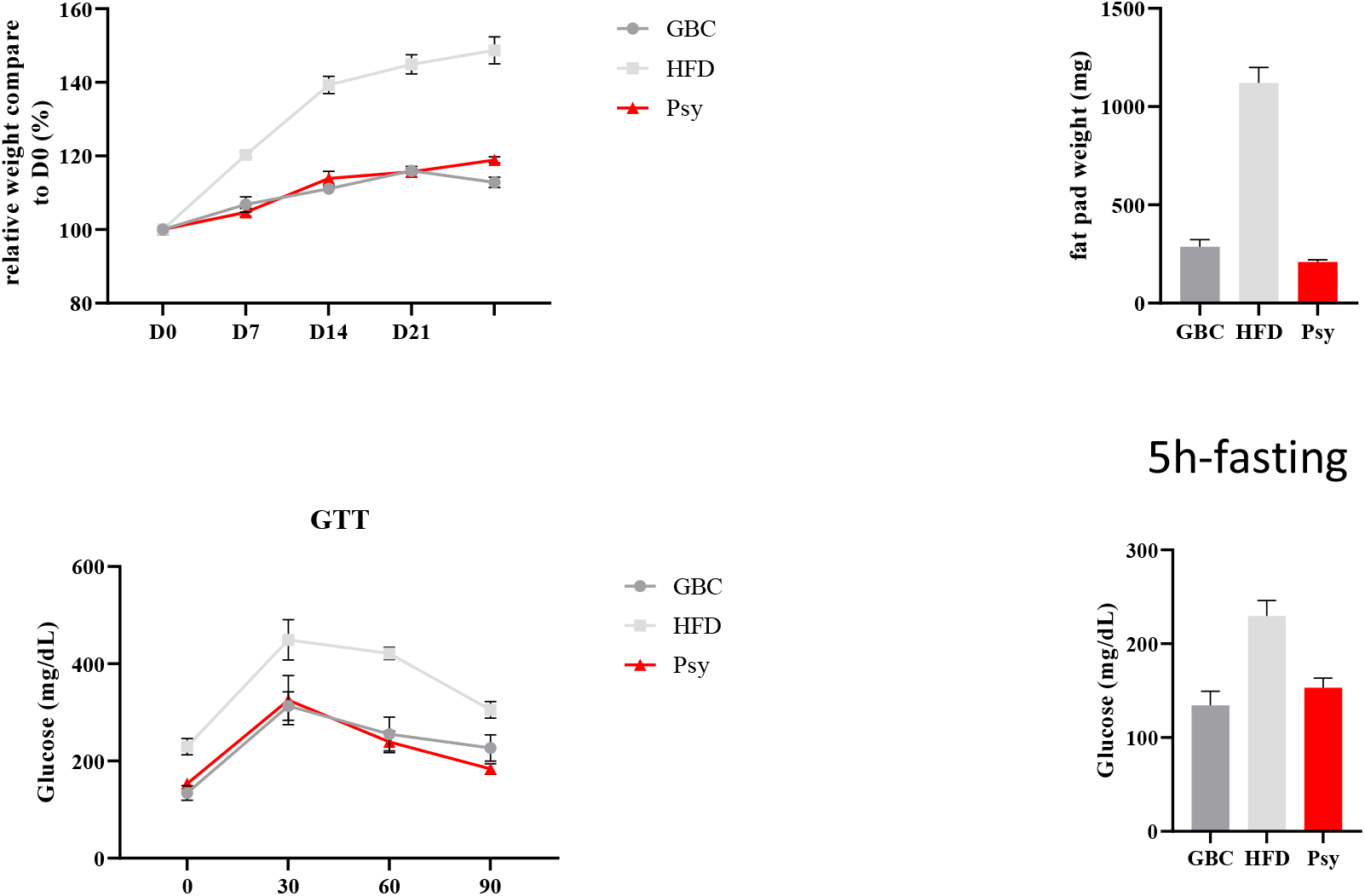
Psyllium protective effect against HFD-induced metabolic syndrome is not dependent of the FXR pathway. Male 6-8 week-old FXRKO male mice were maintained on the specified diet for 7 days and subsequently treated with DSS 2.5% for 9 days. were fed with the indicated diet for 28 day. **A**. Relative body weight over time. **B**. Epididymal fat pad weight. **C**. Glucose tolerance test and respective areas under the curve. **D**. 5 hours fasting glucose level. Data are expressed as means ± SEM of N=4 mice per group. Significance was determined by ANOVA variance test. *P < 0.05, **P < 0.01, ***P < 0.001, ****P < 0.0001.

**Figure 4:**
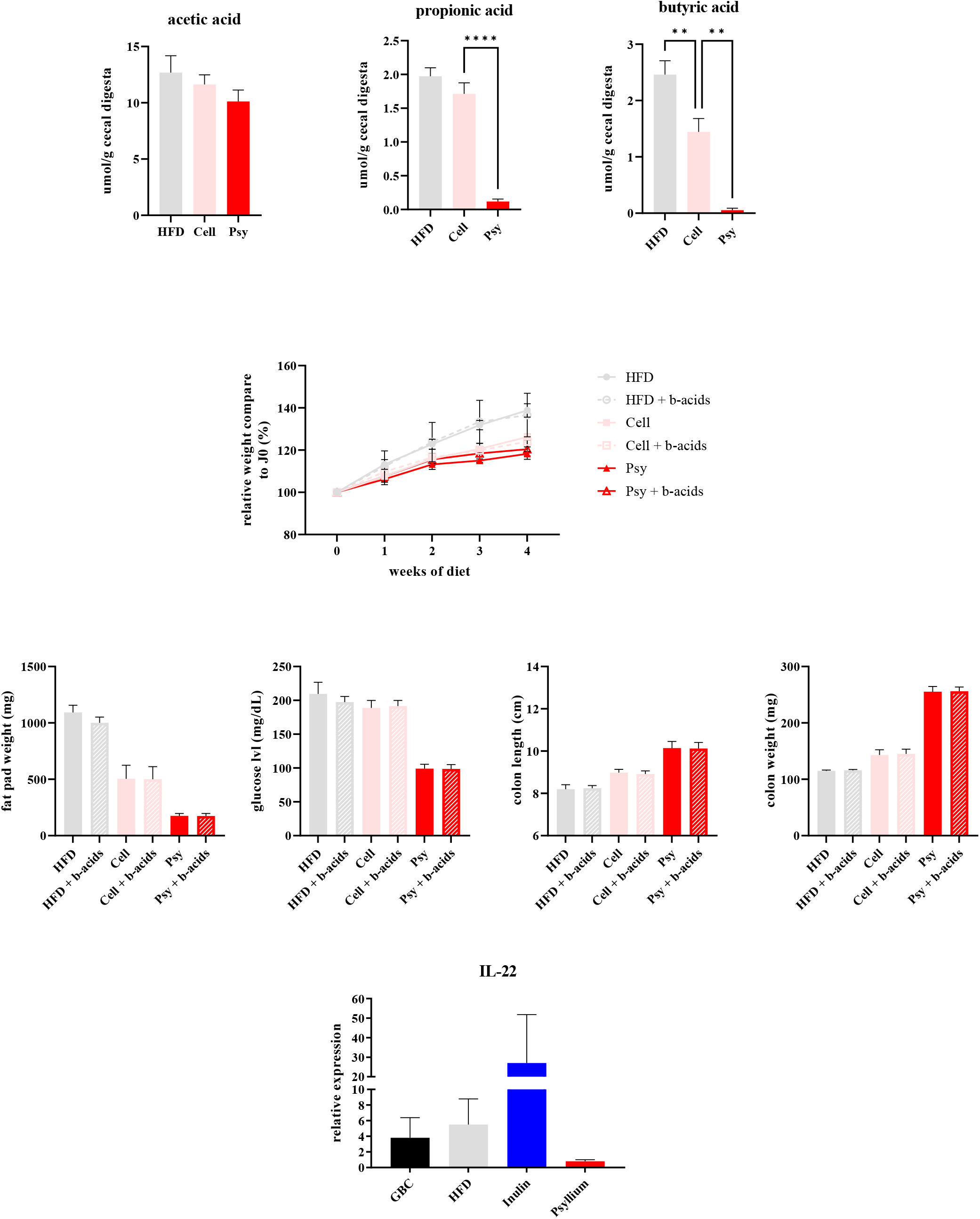
Psyllium did not induce SCFA and did not require fermentation to protect against DSS-induced colitis. Male 6-8 week-old C57Bl/6 mice were fed with the indicated diet for 28 day. **A**. Level of SCFA in the cecum. **B**. Relative body weight over time. **C**. Epididymal fat pad weight. **D**. 5 hours fasting glucose level. **E**. Colon length. **D**. Colon weight. E. Relative expression of IL-22. Data are expressed as means ± SEM of N=5 mice per group. Significance was determined by ANOVA variance test. **P < 0.01, ****P < 0.0001.

**Figure 5:**
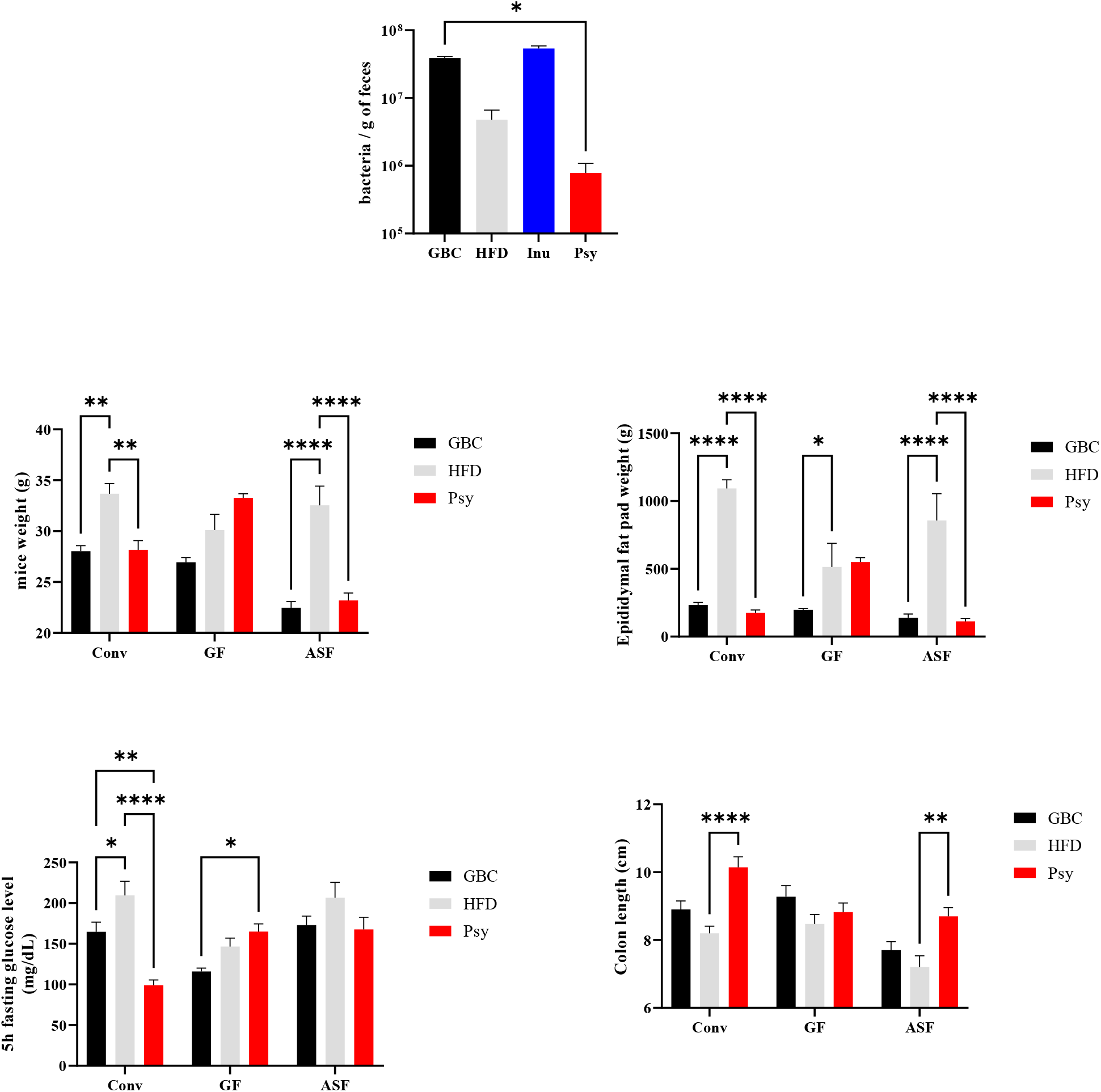
Psyllium induces a drop in bacterial density and required the presence of a microbiota to protect against HFD-induced metabolic syndrome. Male 6-8 week-old C57Bl/6 mice were fed with the indicated diet for 28 day. **A**. Bacterial density in the feces. B-E. Male 6-8 week-old C57Bl/6 conventional, GF and ASF mice were fed with the indicated diet for 28 day. **B**. Relative body weight over time. **C**. Epididymal fat pad weight. **D**. 5 hours fasting glucose level. **E**. Colon length. Data are expressed as means ± SEM of N=5 mice per group for conventional mice, N=x mice per group for the GF mice and N=x mice per group for ASF mice. Significance was determined by ANOVA variance test. *P < 0.05, **P < 0.01 ****P < 0.0001.

Lastly, we sought to discern the role of the microbiota in mediating psyllium’s protection against metabolic syndrome. We observed that complete absence of microbiota, i.e germfree mice, eliminated psyllium’s protection against HFD-induced metabolic syndrome. However, presence of a minimal microbiota, namely an Altered Schaedler Flora, was sufficient to restore psyllium’s ability to protect against metabolic syndrome. Thus, psyllium’s protection requires some microbiota be present for psyllium to protect against diet-induced metabolic syndrome.

## DISCUSSION

Collectively, the results presented herein indicate that enriching an obesogenic diet with psyllium protects against develop of obesity and its associated metabolic abnormalities. Maximal benefits required the diet be comprised of about 8% or more psyllium, which we imagine would be difficult to achieve in humans. Nonetheless, lower doses showed ability ameliorate adiposity. In any case, we expect that better understanding of the mechanism by which psyllium protects against diet-induced metabolic syndrome may yield novel strategies to mitigate this disorder. Psyllium is appreciated to sequester bile acids, thus driving reverse cholesterol transport and potentially impeding lipid uptake. Yet, we would not envision a reason why such actions would require gut microbiota. Thus, how psyllium protects against metabolic syndrome in a microbiota-dependent mechanism is unclear. Future studies are required to better discern the underlying mechanism of its beneficial metabolic impacts.

## Notes

Work supported by NIH grants DK099071 and DK083890 to ATG. AB is supported by CCF RFA 663306.

### Competing Interest Statement

The authors have declared no competing interest.

## References

[1] Zou J, Chassaing B, Singh V, Pellizzon M, Ricci M, Fythe MD, Kumar MV, Gewirtz AT: Fiber-Mediated Nourishment of Gut Microbiota Protects against Diet-Induced Obesity by Restoring IL-22-Mediated Colonic Health. Cell Host Microbe 2018, 23:41–53 e4.

[2] Miles JP, Zou J, Kumar MV, Pellizzon M, Ulman E, Ricci M, Gewirtz AT, Chassaing B: Supplementation of Low- and High-fat Diets with Fermentable Fiber Exacerbates Severity of DSS-induced Acute Colitis. Inflamm Bowel Dis 2017, 23:1133–43.

[3] Bretin A, Zou J, Yeoh BS, Ngo1 VL, Winer S, Winer DA, Reddivari L, Pellizzon M, Walters WA, Patterson AD, Ley R, Chassaing B, Vijay-Kumar M, Gewirtz AT: PSYLLIUM FIBER PROTECTS AGAINST COLITIS VIA ACTIVATION OF BILE ACID SENSOR FXR. CMGH 2023, Under review.

[4] Chassaing B, Gewirtz AT: Mice harboring pathobiont-free microbiota do not develop intestinal inflammation that normally results from an innate immune deficiency. PLoS One 2018, 13:e0195310.

